# A pipelining mechanism supporting previewing during visual exploration and reading

**DOI:** 10.1101/2021.03.25.436919

**Authors:** Ole Jensen, Yali Pan, Steven Frisson, Lin Wang

## Abstract

Humans have a remarkable ability to efficiently explore visual scenes and text by means of eye-movements. Humans typically make eye-movements (saccades) every ~250ms. Since the saccadic motor planning and execution takes 100ms this leaves only ~150ms to recognize the fixated object (or word), while simultaneously previewing candidates for the next saccade goal. We propose a *pipelining mechanism* that efficiently can coordinate visual exploration and reading. The mechanism is timed by alpha oscillations that coordinate the saccades, visual recognition and previewing in the cortical hierarchy. Consequently, the neuronal mechanism supporting visual processing and saccades must be studied in unison to uncover the brain mechanism supporting visual exploration and reading.

**Highlights:**

- Humans have a remarkable ability to efficiently acquire information from visual scenes and pages of text by means of saccadic exploration.
- Visual exploration is surprisingly efficient given the temporal and spatial constraints imposed by the visual system. As such, both information from current fixations as well as upcoming locations must be processed within a 150 ms time window.
- New data recording in humans and non-human primates points to a link between the timing of saccades and alpha oscillations.
- We present a framework in which visual exploration and reading are supported by similar neuronal mechanisms.
- We propose a novel mechanism in which visual exploration and reading is supported by a pipelining mechanism organized by alpha oscillations.
- According to the pipelining mechanism, fixated and previewed objects/words are represented at different phases of an alpha cycle.

## Main text

Our understanding of the visual system presents an intriguing conundrum: How do we manage to efficiently explore visual scenes and text by eye-movements given the relative slow and spatially limited processing capabilities of the human visual system? We saccade every 250 – 300 ms when reading and visually exploring natural scenes. Given that it takes about 100 ms to initiate and execute a saccadic motor program, there is only 150 – 200 ms available to process the fixated object or word while in parallel planning the next saccade. Importantly, since saccades typically land on informative objects or words [1, 2] (Figure 1), a *parafoveal previewing* process is required when exploring and deciding on the next saccade goal. Thus, ultra-fast neuronal computation is essential for supporting saccadic exploration; however, we still need to uncover how the visual system can achieve this remarkable computational feat. The fast computation must rely on a highly tuned machinery in which eye-movements are coordinated with visual input [3]. As such, even small maladaptations can exacerbate problems in visual perception and possibly result in reading disorders.

**Figure 1.**
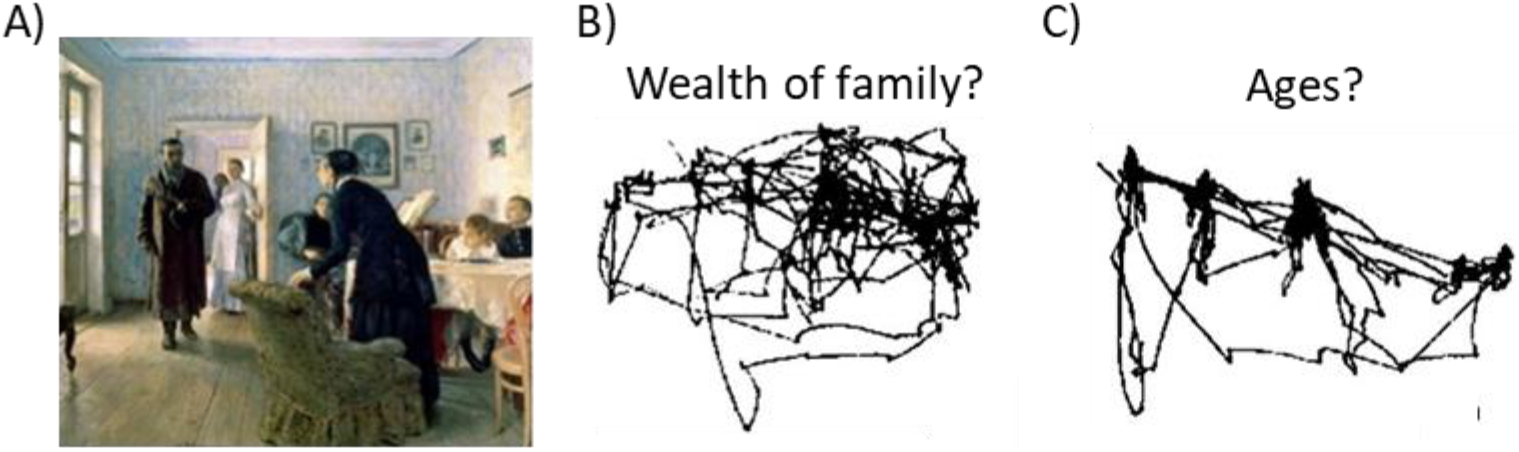
A) When exploring a picture we typically saccade 3-4 times per second. The classic work by Yarbus (1967) demonstrates that saccades land on informative parts of the picture. B) The saccadic path when the participant is asked to judge the wealth of the family. C) The saccadic path when the participant is asked about the ages of the people in the picture. In conclusion, objects explored and thus the saccadic path is depending on the goal of exploration. These findings are consistent with the notion that objects in the scene are previewed prior to saccading to them. Reproduced from Yarbus (1967)

### The temporal constraints during visual exploration and reading

As mentioned above, the identification of current objects (pre-targets) as well as the saccade decision on future saccade goals (targets) must be made within 150 ms after fixating on the current object. This is because saccades are initiated as often as every 250 ms and it takes about 100ms to initiate and execute a saccadic program towards the target [4, 5]. It takes about 60 ms for information to travel from the retina to the visual cortex leaving about 90 ms for neocortical processing of the fixated object (Figure 2A-B). What is the evidence that visual objects can be identified within 150 ms? Multivariate approaches applied to MEG data allow for identifying the *neuronal fingerprint* associated with representation of semantic features [6]. It was found that naturalness and animacy can be decoded from multivariate brain activity at respectively 122 ms and 157 ms after stimulus onset [6]. This timing is consistent with intracranial recordings in monkeys in which object category was decoded within 125 ms in the inferior temporal cortex [7]. Relating visual input to existing memory representations engaged the medial temporal lobe (i.e. parahippocampal areas, entorhinal cortex and hippocampus) at 150 – 200 ms after stimulus onset (for a review see [8]). In humans, event-related potentials increase in response to objects embedded in inconsistent compared to consistent scenes at about 300 – 400 ms (an N400 type response) [9–11]. However, this ERP response is so late that it cannot reflect early recognition. In short, electrophysiological studies suggest that it is possible to identify visual objects at the semantic level (“meaning”) within 150 ms. However, within the same 150 ms time-window future saccade goals must be explored and selected. Given that several processes must be completed within this short time-window, this poses a serious computational challenge to the visual system (Figure 2B). Parafoveal previewing prior to the saccade can serve to reduce recognition of the fixated object to about 110 ms [4, 5, 12]. This then leaves ~40 ms more for previewing the upcoming saccade goal (Figure 2C); i.e. it buys time to alter the saccade plan if, for instance, the saccade goal is deemed uninformative. We suggest that the acceleration of visual processing by previewing might be essential for efficient visual exploration.

**Figure 2.**
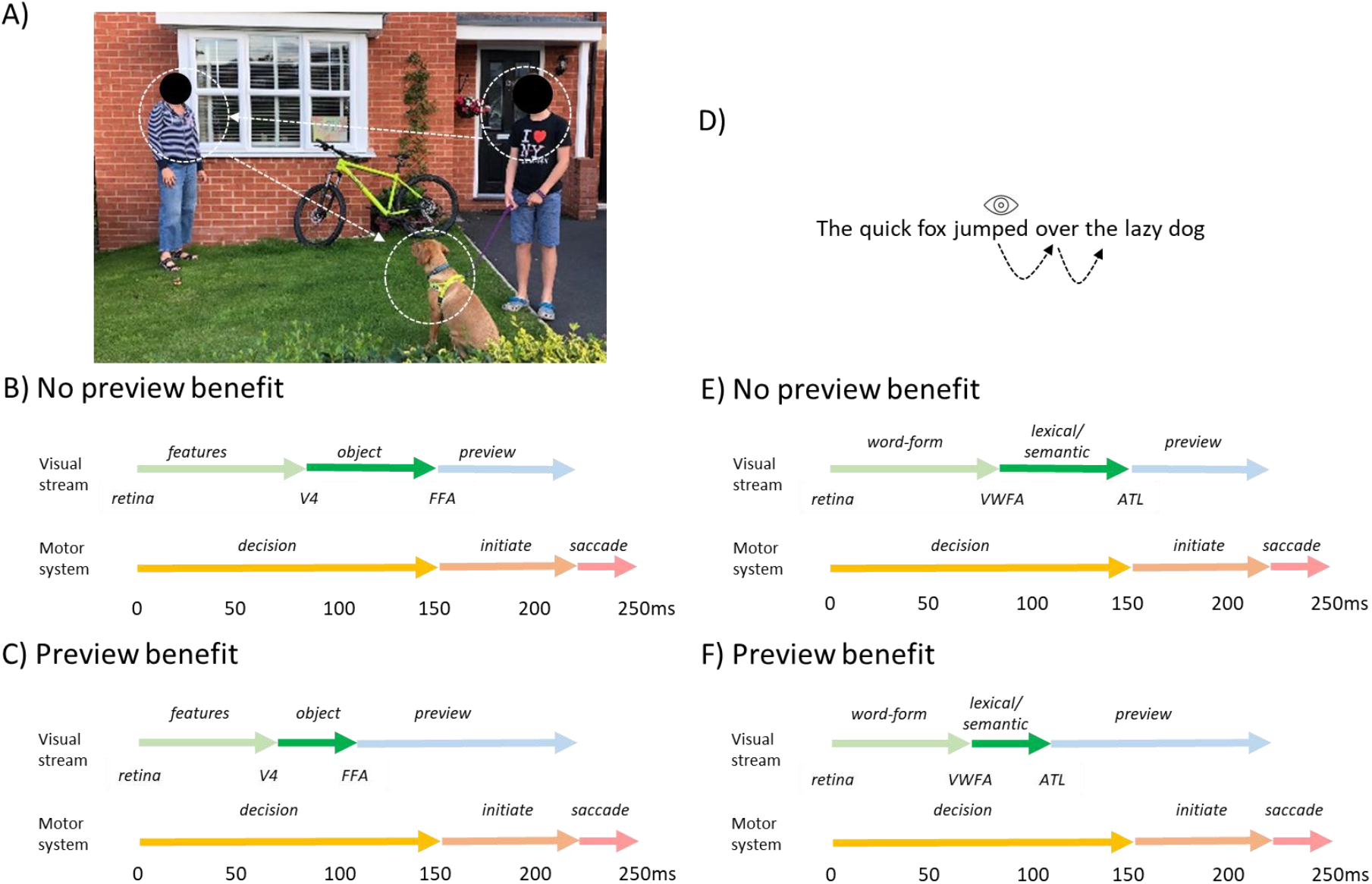
A) In this example the participant first fixates on the *boy* and then saccades to the *woman* followed by the *dog*. B) The timing of the visual exploration process. The visual object (the *boy*) arrives at 60 ms in V1 after which visual features are identified at about 85 ms. There is electrophysiological evidence suggesting that the object can be identified before 150 ms in object selective cortex. While this process is going on, the next saccadic decision must be made (*woman* or *dog?*) such that the motor program can commence. It takes about 100 ms to initiate and execute the saccadic motor command. Typically, a saccade is executed about 250 ms after the fixation onset. As both the object identification as well as the saccade decision must be performed with 150 ms, this places serious computational demands on the visual system. C) Previewing by parafoveal processing allows for speeding up visual processing. For instance, when fixating on the *boy* the *woman* can be previewed. When the *woman* then is fixated the recognition can be reduced to about 110 ms. This has two important advantages: 1) it leaves about 40 ms for previewing the next saccade goal (the *dog*) and 2) the preview occurs sufficiently early to impact the next saccade goal (e.g. to skip). D) A sentence is read by fixating on the words sequentially. When fixating the word *jumped* it must simultaneously be decided on whether to saccade to *over*. E) The timing of the visual reading process. For instance, visual features of the word *over* arrive at 60 ms in V1 after which the word-form is identified in the visual word-form area (VWFA) at ~90 ms. There is electrophysiological evidence suggesting that lexical recognition of the word is done within 150 ms by a network including the anterior temporal lobe (ATL). While this process is going on, the next saccade decision (*over*) must be made such that the saccadic motor program can be initiated. Both the lexical identification as well as the saccade decision must be performed within 150 ms. F) Parafoveal previewing of a word (e.g. *over*) allows for reducing the lexico-semantic identification upon fixation. As such a previewed fixated word could be recognized at 110 ms. This has an important advantage: it leaves more time for previewing the next word and decide the next saccade goal before executing the saccade plan. For instance, a decision might be made to skip highly common but uninformative words.

The temporal constraints during reading are equally tight. After the retinal input has arrived in occipital cortex at 60 ms, the visual word form area (VWFA) is engaged for orthographic processing at about 90 ms [13] (Figure 2D-E). Later follows lexical access and semantic recognition supported by different parts of the left temporal cortex [13]. Eye-tracking research has demonstrated that fixation times are longer for low- compared to high-frequency words [12, 14] and that the word-frequency effect is present a least within 150 ms as revealed by survival analysis [15, 16]. This implies that lexical identification completes before the motor program of the next saccade is initiated [4]; i.e. within 150 ms (Figure 2E). Electrophysiological findings quantifying ERPs and ERFs also support the notion that lexical identification happens within 150 ms [17–19]. As for natural vision, this presents an intriguing problem: if lexical access takes 150 ms and saccade programming about 100 ms, lexical retrieval of the fixated word as well as previewing the next word in the parafovea must be completed within 150 ms [4, 20] (Figure 2E). Previewing will serve to reduce the lexico-semantic recognition possibly to about 110 ms [4, 5, 12], which will leave time to preview the next saccade goal (Figure 2F). This allows just enough time to alter the saccade plan if the next word is deemed uninformative (e.g. *the*).

### The spatial constraints during visual exploration and reading

The visual system also holds an interesting spatial conundrum: Visual acuity drops dramatically for parafoveal vision (2–5 degrees relative to the current fixation) while our eyes can still saccade to relevant (and not necessarily salient) parts of visual scenes [1, 2, 21] (Figure 1). As for visual exploration of natural scenes, it is of great interest to consider the spatial perceptual span in relation to parafoveal visual acuity [22, 23]. Using gaze-contingent paradigms that occlude the peripheral view, it was demonstrated that the radius of the effective visual span is about 8 degrees [24]. Likewise, by applying an *artificial scotoma* that moves with the gaze, participants could still perform a visual search task when occluding up to 4.1 degrees of the field of view. Even peripheral gist information is extracted in this type of paradigm [25]. The relatively large visual span in combination with reduced acuity for parafoveal vision begs the question: in which detail do we process an object before we saccade to it? [26]. In the light of this question, it is debated whether objects are previewed at the semantic level [26, 27]. For instance, studies have shown that search times are faster for objects embedded in inconsistent visual scenes (e.g. a tube of toothpaste in the living room) as compared to consistent scenes (e.g. a tube of toothpaste in the bathroom) [28]; however, it is debated whether this reflects semantic previewing [29, 30]. A recent EEG study investigated fixation-related potentials (FRPs) in response to pre-target objects prior to saccading to target objects that were embedded in either consistent or inconsistent scenes. Larger FRPs were found for objects embedded in inconsistent as compared to consistent scenes [31]. Specifically a larger negative potential at ~300 ms (akin to the N400 type ERP effect) was observed in response to fixations at the pre-target object when the target was inconsistent with the scene. This finding provides support for semantic previewing.

As for reading, the spatial perceptual span in relation to parafoveal visual acuity has been investigated using gaze-contingent paradigms in which performance is measured when the text is occluded to the left and/or right of the gaze. This results in the conclusion that the visual span extends 14–15 letter spaces (2–3 words) to the right of fixation and 3–4 letters to the left [32]. Interestingly, this effect is reversed in readers of Hebrew who read from right to left [33]. Therefore, the spatial perceptual span in reading is also not constrained by the reduction in visual acuity of extra-foveal vision. However, it has been shown that occluding the word just to the right of fixation, reduces reading-speed by 25–40 ms per word [12]. Interestingly, making the fixated word disappear after 60 ms hardly impacts reading, but making the parafoveal word disappear after 60ms, increases reading times substantially [34]. As such there is strong evidence that fluent reading relies on previewing.

Again, in which detail is upcoming text previewed in the parafovea? There is strong evidence for previewing at the sub-lexical level (e.g. orthographic and phonological levels). Experiments using the gaze-contingent boundary paradigm demonstrated that fixation times on a word were reduced after it was primed in the parafovea by an orthographically similar letter string (e.g *sorp* priming *song*) compared to an unrelated condition (*door* replaced by *song*) (e.g [35–37]). A similar effect was found with respect to phonological previewing using homophones (e.g., *beach* priming *beech*) [38, 39]. Previewing at the phonological level is also supported by reading studies manipulating the spelling-sound regularity. It is well established that fixation times are longer on words with irregular spelling-sound mappings. However, this difference disappears when previewing is prevented in a gaze contingent paradigm [40].

It remains debated if words are previewed at the lexical and semantic level. Previewing at the *lexical level* has been investigated using sentences containing target words of low- or high-lexical frequency. Several eye-tracking studies have found that pre-target fixation times are not modulated by the lexical frequency of the target word, suggesting the absence of lexical previewing [41]. We recently challenged this notion by combining eye-tracking with a rapid frequency tagging paradigm. In this paradigm, we subliminally flickered the target words that were of low- or high-frequency during natural reading. We found that when readers fixated on the pre-target words, there was a stronger tagging response as measured by MEG when the target words were of low- compared to high-frequency [42]. This finding provides support for previewing at the lexical level. Another controversial topic is whether there is previewing at the *semantic level*. This question was studied using boundary paradigms in which the target word is changed just as the participant saccades to it. When for instance the target word changed from *tune* to *song,* the fixation times on *song* were compared to when the target word changed from the unrelated word *door* to *song*. The fixation times on *song* did not differ between these two conditions when the sentences were presented in English, indicating a lack of semantic preview benefits [43]. Surprisingly, studies using Chinese [44] and German [45] have found evidence for semantic previewing, e.g. shorter fixation times on *dog* when the previewed word was *puppy* compared to when it was *desk* (for the German study, this was possibly explained by the capitalization of nouns [46]). Another EEG study was based on participants reading lists of nouns that were either semantically related or not. In favour of no semantic previewing [47], the fixation-related potentials for a given word did not depend on whether the preceding word was semantically related to it or not. In sum, while there is evidence for previewing at the sub-lexical level, there are mixed reports on lexical and semantic previewing.

### A mechanism supporting visual exploration and reading by pipelining coordinated by alpha oscillations

Having introduced the temporal and spatial constraints of the visual system, we here propose a pipelining mechanism that can be used to guide efficient visual exploration and reading. Before explaining the details of the mechanism, we will first review the temporal coding scheme observed in exploring animals that inspired the model. There is an intriguing link between visual and spatial exploration. The goal of both behaviours is to process information from the current location while deciding where to go next. Intracranial neuronal investigations in behaving rats have provided important insights into the neurophysiological mechanisms coordinating this process. Place cell recordings in the rat hippocampus have demonstrated that neuronal theta oscillations (6 – 12 Hz) play an essential role for organizing neuronal representations of space. The phenomenon of *theta phase precession* shows that a given place cell fires late in the theta cycle as the rat enters a place field. As the rat advances, the firing precesses to earlier theta phases. This finding is best explained by a mechanism in which a sequence of spatial representations is ‘read out’ within a theta cycle [48, 49]. Neuronal representations early in the theta cycle code for current location whereas firing later in the cycle codes for upcoming locations. This phase-coding scheme is consistent with a pipelining mechanism in which different representations along the path are sequentially processed at different theta phases. Could a related mechanism support visual exploration in which objects on a *saccadic path* are encoded as a sequence organized by oscillations?

In support of the above scheme, phase-coding with respect to neuronal oscillations has been identified from intracranial recordings in humans performing visual and working memory tasks. For instance, using human intracranial data it was demonstrated that individual working memory representations are represented at different phases of an 8 Hz rhythm [50]. Another intracranial study found that different visual categories were reflected by a phase-code [51]. Work based on intracranial recordings in non-human primates also reports phase coding in the visual system in various tasks [52–54]. Inspired by the phase-coding mechanism we propose a framework (Box 1 and 2) where oscillations in the 8–13 Hz alpha band serve to organize visual presentations in a phase-coded manner to support parafoveal previewing and eventually guide the saccadic trajectory.

We hypothesize that natural visual exploration and reading relies on a process in which several objects or words are processed simultaneously at different levels in the cortical hierarchy. Consider Box 1A in which the viewer fixates on the *woman.* The visual input propagates in the cortical hierarchy in which features of increasing complexity are combined to eventually recognize the object *woman* in inferior temporal (IT) cortex. While the participant fixates on the *woman*, the *dog* is previewed as a potential saccade goal. The previewing creates a bottleneck problem in IT cortex when two objects (e.g. *woman* and *dog*) are processed. We propose that the bottleneck-problem is solved by a pipelining mechanism in which several objects are processed simultaneously but at different levels in the cortical hierarchy (detailed in Box 1B). This scheme serves two different purposes: First, the preview of the *dog* will speed up the visual processing when this object eventually is fixated, thus reducing the recognition time. It is essential that the saccades are locked to the phase of the alpha oscillations in order for the processing to be coordinated. This scheme increases the efficacy of visual processing and it also buys some time allowing for the saccade plan to be revised in case the previewed object is deemed irrelevant.

We argue that a similar mechanism supports natural reading (see Box 2), with the exception that the saccades typically are directed to the right. When the word *jumped* is fixated, this allows for *over* to be previewed. To reduce a bottle-neck problem when processing the words, they are organized in a temporal coding manner along the alpha cycle. Parafoveal previewing results in the lexico-semantic identification reducing from about 150 ms to 110 ms. This provides extra time for evaluating the next saccade goal and potentially skip a less informative word (e.g. *the*). Importantly, the saccades are locked to the phase of the alpha oscillations in order to organize the timing of the processing and the visual input. In short, we argue that efficient visual exploration and reading rely on parafoveal previewing, and the created bottleneck problem can be solved by a pipelining mechanism, suggesting that the processing of fixated and previewed objects is coordinated in time by alpha oscillations.

A computational mechanism organized as a pipeline requires an intricate temporal organization (Box 1 and 2). The transfer of representations between levels in the hierarchy as well as the sequential processing must be coordinated. In the example Box 1, some of the visual features of the *boy* will propagate to face-selective areas. Likewise, the face-selective area will process the *boy* slightly earlier than the *woman*. Based on recent findings, we propose that the oscillatory coupling serves to coordinate the information transfer between regions [55, 56] as well as organizing the sequential processing in a phase-coded manner.

Which neuronal dynamics might support a pipelining mechanism coordinated by brain oscillations? Based on human and animal data, it has been proposed that theta and alpha oscillations are a consequence of pulses of inhibition [57–59]. At the peak of an inhibitory pulse, neurons are prevented from firing. As the inhibition ramps down over the cycle, the most excitable neurons will fire first, then the somewhat less excited neurons and so forth. As such, the pulses of inhibition implement a type of filter, ensuring that neuronal representations are activated sequentially according to excitability [59, 60]. This mechanism can account for the theta phase precession in rats [57] and has been proposed to operate in the visual system [59]. In case of the visual system, fixated representations are more excited compared to parafoveal representations. This allows the foveal representation to overcome the inhibition earlier and thus activate earlier in the alpha cycle. The sorting of visual representations, according to excitability, is a crucial component in the proposed pipelining mechanism (Box 1 and 2). While we have put forward an example with 2 objects in each cycle, the scheme could be extended to 3-4 objects and to include more hierarchical levels.

Computational modelling has proposed mechanism for how phase-coded representations are exchanged between brain regions [55]. Indeed, intracranial recordings in non-human primates suggest that synchronization in the alpha band reflects forward communication in the extended visual system [61]. The phase of the alpha oscillations modulates the gamma band activity, which serves to segment the representations in the alpha cycle. This scheme allows for the exchange of phase-coded representations between brain regions [62]. We propose that alpha-band phase-synchronization in the cortical hierarchy supports the phase-coded pipelining scheme by coordinating the feedforward transfer of increasingly higher level representations.

### Evidence linking alpha oscillations and saccades

As outlined in Box 1 and 2, we propose that alpha oscillations are involved in organizing a pipelining mechanism supporting visual processing. While alpha oscillations for decades were thought to reflect idling or a state of rest [63], it is now evident that they are involved in numerous cognitive processes [58, 64, 65]. One key observation is that alpha oscillations are present during continuous visual presentation e.g. [66, 67]. More recently, both human and animal research has found an intriguing link between the phase of alpha oscillations and saccades. A study based on both MEG and intracranial human data showed that saccade onsets are locked to the phase of ongoing alpha oscillations when viewing natural images [68]. Importantly, the locking of saccades to alpha phase was stronger for pictures later remembered as compared to later forgotten. This suggests that visual information impact memory areas stronger when saccades are coordinated by the phase of alpha oscillations. This notion is further supported by a nonhuman primate study in which saccades produced a phase-reset of theta/alpha oscillations in the hippocampus; the magnitude of the phase-reset predicted memory encoding [69]. Using EEG, it was shown that saccadic reaction times relates to the phase of pre-saccadic alpha oscillations [70]. A recent study in non-human primates reported on multi-electrode recording in the frontal-eye-field that allowed for decoding of the focus of the attentional spotlight. Importantly, the spotlight explored the visual space at a 7-12 Hz alpha rhythm (‘attentional saccades’) [71]. In another non-human primate study, signals from the V4 receptive field of respectively current and future fixations were coherent in the alpha band around the time of saccades [72]. Finally, saccadic suppression is also related to alpha oscillations: recording sites in V4 associated with peripheral vision increase in alpha power during saccades [73]. In short, strong evidence is accumulating in support of an intimate connection between saccades and alpha oscillations. These findings in humans and animals provide some support for the proposed mechanism for visual exploration and reading (Box 1 and 2)

### Previewing primes and speeds up visual recognition

Key to the mechanism that we propose is that previewing speeds up the identification of objects in visual scenes as well as lexico-semantic retrieval during reading [42, 74, 75]. Several models suggest that object and lexical recognition relies on attractor type networks (e.g. [76–79]). The core idea is that incomplete representations can be reconstructed through pattern completion in a network of neurons with recurrent connections. The time it takes for an attractor network to converge is dependent on the basin of attraction and the trajectory to be travelled. *Priming* can be thought of as guiding a neuronal representation closer to the attractor as studied in computational models [80]. We propose that parafoveal previewing serves to prime a given neuronal representation so that when the respective object or word is fixated, retrieval is facilitated, and identification is sped up. In short, attractor type networks can provide a physiologically compatible account for how parafoveal previewing can prime object identification and lexico-semantic retrieval.

### Concluding remarks

We argue that the visual system must operate in a highly efficient manner to support visual exploration and reading. The core issue is that the fixated object or word must be processed in the same time-interval as when the next saccade goal is planned. Given the bottleneck problem in the visual hierarchy, we propose that this is achieved by a pipelining mechanism in which current and upcoming spatial locations are processed – not simultaneously – but in fast succession. Importantly, we propose that neuronal oscillations in the alpha band serve to coordinate the pipelining mechanism that is implemented by a phase-coding scheme in which different representations activate sequentially along the phase of the alpha oscillations. Finally, to time the visual input with the neuronal processing, the mechanism also requires that saccades are locked to the phase of the alpha oscillations.

The framework results in a set of testable predictions. A core prediction is that alpha band oscillations coordinate the neuronal processing associated with saccadic visual exploration. As a result, representationally-specific representations for fixated and upcoming saccade goals should be coupled to the phase of the alpha oscillations. This could be tested by MEG or EEG recordings combined with eye-tracking studies in humans engaged in visual exploration or reading tasks. The time-course of the representationally specific activation could be identified by multivariate approaches and related to the phase of the ongoing alpha oscillations [81]. As a complementary approach, rapid frequency tagging at different frequencies (50 to 70 Hz) could be used to track several objects and the respective neuronal signals would then reflect the neuronal processing [42]. Specifically, we predict that the fixated as well as the parafoveal object (or word) would be become active at different phases of the alpha signal (see Box1C and Box2B). Importantly, we also predict that saccades would be locked to alpha phase (as in [68]) and that this locking would be more pronounced with an increase in task-demands. Finally, we predict that regions in the cortical hierarchy are phase-locked in the alpha band. The phase-locking would allow for the phase-coded information to be exchanged [55, 56]; possibly there would be a systematic phase-shift of the alpha cycles through the cortical hierarchy to guide the feed-forward flow of phase-coded information [82].

In sum, we have here presented a novel and testable framework for the neuronal mechanisms supporting visual exploration and reading in relation to saccades. Crucially, neuronal oscillations are required for organizing the visual representation as well as the timing of saccades. Since the proposed mechanism provides a unified account for visual exploration and reading, it also opens the door for future investigations aimed at understanding the neuronal substrate associated with reading disorders.

## Outstanding Questions

In which detail are upcoming objects previewed before saccades are made to them? Are they previewed at the semantic level or maybe just in terms of features?

- In which detail are upcoming words previewed during reading before saccades are made to them? Are they primarily previewed at the sub-lexical (e.g. orthographical, phonological, orthographic) or also at the lexical and semantic level?
- What is the role of brain oscillations in visual exploration and reading? Recent studies have found cases where saccades are locked to the phase of alpha oscillations [68], but how general is this phenomenon?
- Are different objects and words represented along the phase of oscillations in the alpha cycle during visual exploration and reading (akin to the coding scheme of place representation organized by theta oscillations observed in exploration rats)? This can be addressed using multi-variate methods applied to MEG and EEG data in order to relate representational specific information to the phase of oscillations in the alpha band [81]. Likewise, *rapid frequency tagging* can be used to investigate the allocation of visual resources already prior to a saccade in relation to brain oscillations [42].
- Is there a link between previewing abilities and reading disorders? For instance, impaired previewing during reading might account for some of the reading deficits observed in some types of developmental and acquired dyslexia. Can our proposed pipelining mechanism account for the impaired previewing?
- Recent evidence suggests that prediction plays an important role in language comprehension [83–85]. How do top-down predictions impact bottom-up parafoveal previewing during reading? Could it be that both pre-activated representations and the representations of the previewed objects are encoded at the same phase of the alpha cycle?

**Box 1:**
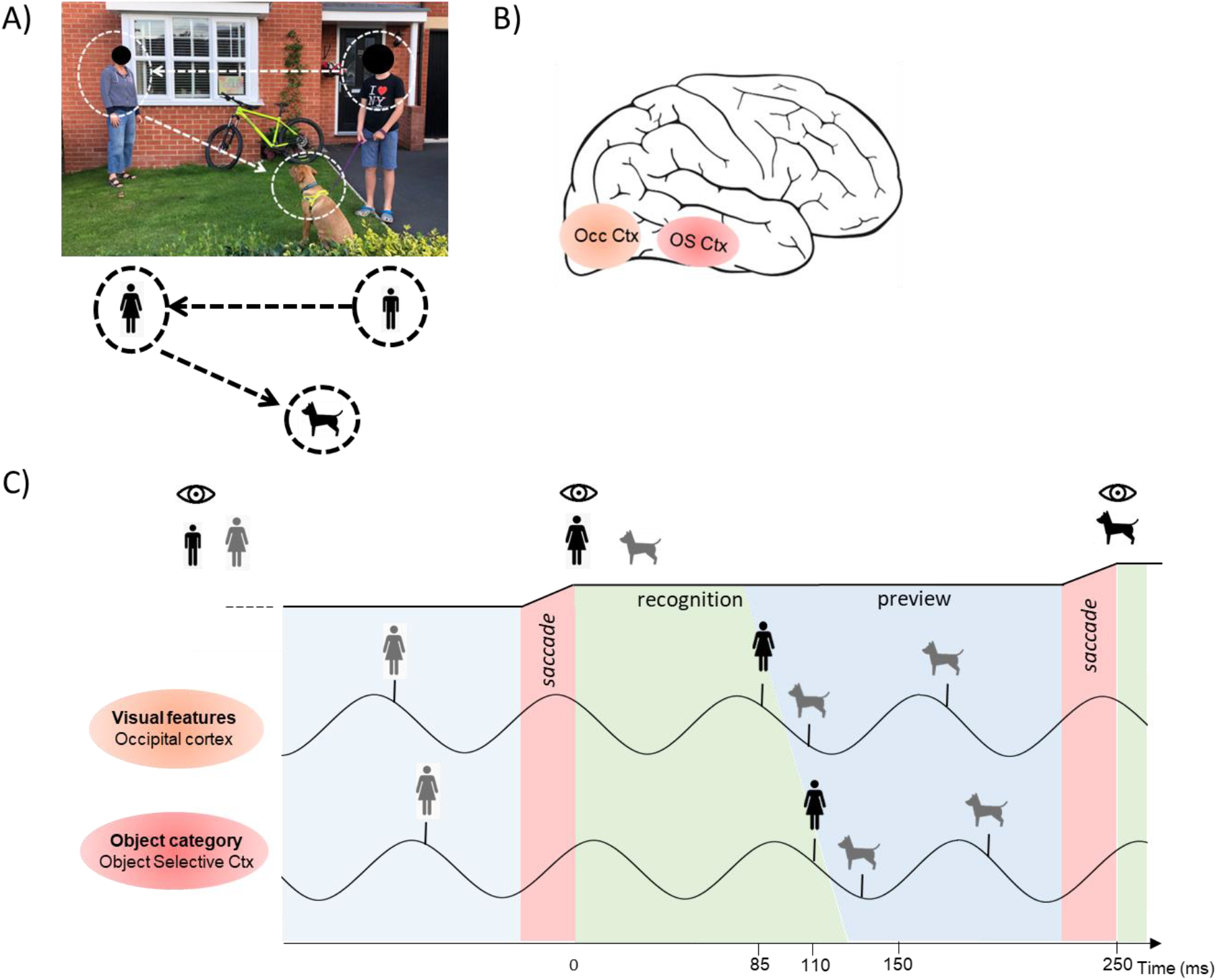
A model for pipelining during visual exploration. A) The participant fixates on the *boy* and then saccades to the *woman* and finally to the *dog*. B) For simplicity, we assume two stages in which simple features (e.g. color) are identified first, followed by object category recognition in respectively V4 in occipital cortex and objects selective cortex in the inferior temporal lobe. C) We hypothesize that the temporal organization supporting the pipelining mechanism is coordinated by oscillations in the alpha band. In this example, 12 Hz alpha oscillations can be considered pulses of inhibition repeated every 83 ms. At the peak of the alpha cycle, neuronal firing is inhibited. As the inhibition ramps down the most excitable representation will activate and so forth. Consider time point t = 0 ms in which the participant moves the eyes from the *boy* and fixates on the *woman*. We assume that saccades are locked to the phase of the alpha oscillations such that the visual input of the *woman* arrives at the early down-going inhibitory slope of the alpha cycle at about t = 85 ms where simple visual features of the woman engage visual occipital cortex (e.g. colors in V4). These feature representations are projected to object selective cortex for category identification by 110 ms. This fast category identification is made possible by the preview of the *woman* prior to the saccade which has primed the ‘semantic’ access. Importantly, the pipelining scheme allows the *woman* and the *dog* to be processed in the same cycle in a multiplex manner thus avoiding bottleneck problems. This scheme allows for a fast decision to be made to either saccade to the *dog* or hold the saccade and preview another object as a potential target.

**Box 2:**
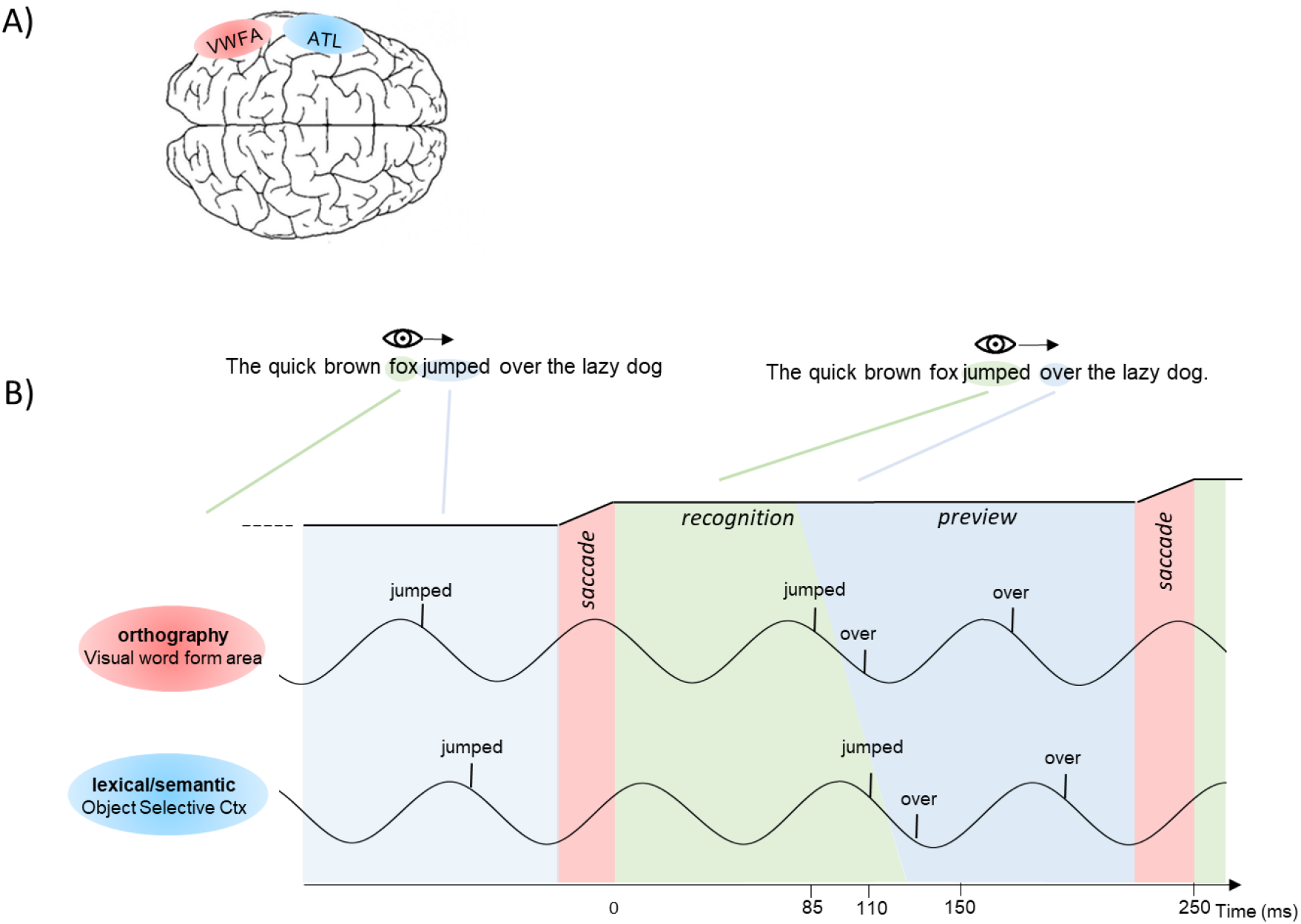
A model for pipelining during natural reading. A) For the sake of simplicity, two stages of word recognition are assumed, namely orthographic identification in visual word from area (VWFA) followed by lexico-semantic access in an extended network including the anterior temporal lobe. B) We hypothesize that the temporal organization supporting the pipelining mechanism is coordinated by oscillations in the alpha band. The 12 Hz alpha oscillations can be considered pulses of inhibition repeated every 83 ms [58]. Consider time point t = 0 ms in which the subject saccades and fixates on *jumped*. We assume that saccades are locked to the phase of the alpha oscillations such that the visual input of *jumped* arrives at the early down-going inhibitory slope of the alpha cycle at about 85 ms for orthographic feature identification in the VWFA. The orthographic representations propagate to the ATL for lexico-semantic identification by 110 ms. This fast lexico-semantic process is made possible by the preview of *jumped* prior to the saccade which has primed the lexical access. Importantly, the pipeline scheme allows both *jumped* and *over* to be lexico-semantically processed in the same cycle but at slightly different points in time, thus avoiding a bottleneck problems in the ATL. During the fixation of *over* the word *the* is previewed. Given that the word *the* carries little information, a decision to skip can be made. However, this is only possible if *over* has been previewed since this will speed up the processing of *over* leaving more time to preview *the*.

## References

1. Yarbus, A.L. (1967) Eye movements and vision, Plenum Press.

2. Torralba, A. et al. (2006) Contextual guidance of eye movements and attention in real-world scenes: the role of global features in object search. Psychol Rev 113 (4), 766–86.

3. Liversedge, S.P. and Findlay, J.M. (2000) Saccadic eye movements and cognition. Trends Cogn Sci 4 (1), 6–14.

4. Reichle, E.D. and Reingold, E.M. (2013) Neurophysiological constraints on the eye-mind link. Front Hum Neurosci 7, 361.

5. Becker, W. and Jurgens, R. (1979) An analysis of the saccadic system by means of double step stimuli. Vision Res 19 (9), 967–83.

6. Cichy, R.M. et al. (2014) Resolving human object recognition in space and time. Nat Neurosci 17 (3), 455–62.

7. Hung, C.P. et al. (2005) Fast readout of object identity from macaque inferior temporal cortex. Science 310 (5749), 863–6.

8. Mormann, F. et al. (2008) Latency and selectivity of single neurons indicate hierarchical processing in the human medial temporal lobe. J Neurosci 28 (36), 8865–72.

9. Ganis, G. and Kutas, M. (2003) An electrophysiological study of scene effects on object identification. Brain Res Cogn Brain Res 16 (2), 123–44.

10. Mudrik, L. et al. (2010) ERP evidence for context congruity effects during simultaneous object-scene processing. Neuropsychologia 48 (2), 507–17.

11. Vo, M.L. and Wolfe, J.M. (2013) Differential electrophysiological signatures of semantic and syntactic scene processing. Psychol Sci 24 (9), 1816–23.

12. Rayner, K. (1998) Eye movements in reading and information processing: 20 years of research. Psychol Bull 124 (3), 372–422.

13. Carreiras, M. et al. (2014) The what, when, where, and how of visual word recognition. Trends Cogn Sci 18 (2), 90–8.

14. Rayner, K. (2009) Eye Movements in Reading: Models and Data. J Eye Mov Res 2 (5), 1–10.

15. Reingold, E.M. et al. (2012) Direct lexical control of eye movements in reading: evidence from a survival analysis of fixation durations. Cogn Psychol 65 (2), 177–206.

16. Sheridan, H. and Reichle, E.D. (2016) An Analysis of the Time Course of Lexical Processing During Reading. Cogn Sci 40 (3), 522–53.

17. Sereno, S.C. et al. (2003) Context effects in word recognition: evidence for early interactive processing. Psychol Sci 14 (4), 328–33.

18. Assadollahi, R. and Pulvermuller, F. (2003) Early influences of word length and frequency: a group study using MEG. Neuroreport 14 (8), 1183–7.

19. Dambacher, M. et al. (2006) Frequency and predictability effects on event-related potentials during reading. Brain Res 1084 (1), 89–103.

20. Sereno, S.C. and Rayner, K. (2000) The when and where of reading in the brain. Brain and Cognition 42 (1), 78–81.

21. Liversedge, S.P. et al. (2013) The Oxford handbook of eye movements, 2013 edition edn., Oxford University Press.

22. Nuthmann, A. and Henderson, J.M. (2010) Object-based attentional selection in scene viewing. J Vis 10 (8), 20.

23. Pajak, M. and Nuthmann, A. (2013) Object-based saccadic selection during scene perception: evidence from viewing position effects. J Vis 13 (5).

24. Nuthmann, A. (2013) On the visual span during object search in real-world scenes. Visual Cognition 21 (7), 803–837.

25. Larson, A.M. and Loschky, L.C. (2009) The contributions of central versus peripheral vision to scene gist recognition. J Vis 9 (10), 6 1–16.

26. Wu, C.C. et al. (2014) Guidance of visual attention by semantic information in real-world scenes. Front Psychol 5, 54.

27. Treisman, A.M. and Gelade, G. (1980) Feature-Integration Theory of Attention. Cognitive Psychology 12 (1), 97–136.

28. LaPointe, M.R.P. and Milliken, B. (2016) Semantically incongruent objects attract eye gaze when viewing scenes for change. Visual Cognition 24 (1), 63–77.

29. Vo, M.L.H. and Henderson, J.M. (2011) Object-scene inconsistencies do not capture gaze: evidence from the flash-preview moving-window paradigm. Attention Perception & Psychophysics 73 (6), 1742–1753.

30. Underwood, G. (2009) Cognitive Processes in Eye Guidance: Algorithms for Attention in Image Processing. Cognitive Computation 1 (1), 64–76.

31. Coco, M.I. et al. (2020) Fixation-related Brain Potentials during Semantic Integration of Object-Scene Information. Journal of Cognitive Neuroscience 32 (4), 571–589.

32. Rayner, K. et al. (1980) Asymmetry of the effective visual field in reading. Percept Psychophys 27 (6), 537–44.

33. Pollatsek, A. et al. (1981) Asymmetries in the perceptual span for Israeli readers. Brain Lang 14 (1), 174–80.

34. Rayner, K. et al. (2006) Eye movements when reading disappearing text: the importance of the word to the right of fixation. Vision Res 46 (3), 310–23.

35. Rayner, K. et al. (1980) Integrating information across eye movements. Cogn Psychol 12 (2), 206–26.

36. McConkie, G.W. and Zola, D. (1979) Is visual information integrated across successive fixations in reading? Percept Psychophys 25 (3), 221–4.

37. Rayner, K. et al. (1986) Against parafoveal semantic preprocessing during eye fixations in reading. Can J Psychol 40 (4), 473–83.

38. Chace, K.H. et al. (2005) Eye movements and phonological parafoveal preview: effects of reading skill. Can J Exp Psychol 59 (3), 209–17.

39. Pollatsek, A. et al. (1992) Phonological codes are used in integrating information across saccades in word identification and reading. J Exp Psychol Hum Percept Perform 18 (1), 148–62.

40. Sereno, S.C. and Rayner, K. (2000) Spelling-sound regularity effects on eye fixations in reading. Percept Psychophys 62 (2), 402–9.

41. Degno, F. et al. (2019) Parafoveal previews and lexical frequency in natural reading: Evidence from eye movements and fixation-related potentials. J Exp Psychol Gen 148 (3), 453–474.

42. Pan, Y.F. S.,; Jensen, O. (2020) Lexical parafoveal processing in natural reading predicts reading speed. bioRxiv.

43. Rayner, K. et al. (2003) On the processing of meaning from parafoveal vision during eye fixations in reading. Mind’s Eye: Cognitive and Applied Aspects of Eye Movement Research, 213–234.

44. Yan, M. et al. (2009) Readers of Chinese extract semantic information from parafoveal words. Psychon Bull Rev 16 (3), 561–6.

45. Hohenstein, S. et al. (2010) Semantic preview benefit in eye movements during reading: A parafoveal fast-priming study. J Exp Psychol Learn Mem Cogn 36 (5), 1150–70.

46. Rayner, K. and Schotter, E.R. (2014) Semantic preview benefit in reading English: The effect of initial letter capitalization. J Exp Psychol Hum Percept Perform 40 (4), 1617–28.

47. Dimigen, O. et al. (2012) Trans-saccadic parafoveal preview benefits in fluent reading: A study with fixation-related brain potentials. Neuroimage 62 (1), 381–393.

48. Skaggs, W.E. et al. (1996) Theta phase precession in hippocampal neuronal populations and the compression of temporal sequences. Hippocampus 6 (2), 149–72.

49. Jensen, O. and Lisman, J.E. (1996) Hippocampal CA3 region predicts memory sequences: accounting for the phase precession of place cells. Learn Mem 3 (2-3), 279–87.

50. Bahramisharif, A. et al. (2018) Serial representation of items during working memory maintenance at letter-selective cortical sites. PLoS Biol 16 (8), e2003805.

51. Watrous, A.J. et al. (2015) Phase-amplitude coupling supports phase coding in human ECoG. Elife 4.

52. Kayser, C. et al. (2012) Analysis of slow (theta) oscillations as a potential temporal reference frame for information coding in sensory cortices. PLoS Comput Biol 8 (10), e1002717.

53. Montemurro, M.A. et al. (2008) Phase-of-firing coding of natural visual stimuli in primary visual cortex. Curr Biol 18 (5), 375–80.

54. Turesson, H.K. et al. (2012) Category-selective phase coding in the superior temporal sulcus. Proc Natl Acad Sci U S A 109 (47), 19438–43.

55. Jensen, O. (2001) Information transfer between rhythmically coupled networks: reading the hippocampal phase code. Neural Comput 13 (12), 2743–61.

56. Bonnefond, M. et al. (2017) Communication between Brain Areas Based on Nested Oscillations. eNeuro 4 (2).

57. Mehta, M.R. et al. (2002) Role of experience and oscillations in transforming a rate code into a temporal code. Nature 417 (6890), 741–6.

58. Mazaheri, A. and Jensen, O. (2010) Rhythmic pulsing: linking ongoing brain activity with evoked responses. Front Hum Neurosci 4, 177.

59. Jensen, O. et al. (2014) Temporal coding organized by coupled alpha and gamma oscillations prioritize visual processing. Trends Neurosci 37 (7), 357–69.

60. Gips, B. et al. (2016) A biologically plausible mechanism for neuronal coding organized by the phase of alpha oscillations. Eur J Neurosci 44 (4), 2147–61.

61. Saalmann, Y.B. et al. (2012) The pulvinar regulates information transmission between cortical areas based on attention demands. Science 337 (6095), 753–6.

62. Quax, S. et al. (2017) Top-down control of cortical gamma-band communication via pulvinar induced phase shifts in the alpha rhythm. PLoS Comput Biol 13 (5), e1005519.

63. Pfurtscheller, G. et al. (1996) Event-related synchronization (ERS) in the alpha band--an electrophysiological correlate of cortical idling: a review. Int J Psychophysiol 24 (1-2), 39–46.

64. Van Diepen, R.M. et al. (2019) The functional role of alpha-band activity in attentional processing: the current zeitgeist and future outlook. Curr Opin Psychol 29, 229–238.

65. Palva, S. and Palva, J.M. (2007) New vistas for alpha-frequency band oscillations. Trends Neurosci 30 (4), 150–8.

66. Wang, L. et al. (2018) Language Prediction Is Reflected by Coupling between Frontal Gamma and Posterior Alpha Oscillations. J Cogn Neurosci 30 (3), 432–447.

67. VanRullen, R. and Macdonald, J.S. (2012) Perceptual echoes at 10 Hz in the human brain. Curr Biol 22 (11), 995–9.

68. Staudigl, T. et al. (2017) Saccades are phase-locked to alpha oscillations in the occipital and medial temporal lobe during successful memory encoding. PLoS Biol 15 (12), e2003404.

69. Jutras, M.J. et al. (2013) Oscillatory activity in the monkey hippocampus during visual exploration and memory formation. Proc Natl Acad Sci U S A 110 (32), 13144–9.

70. Drewes, J. and VanRullen, R. (2011) This is the rhythm of your eyes: the phase of ongoing electroencephalogram oscillations modulates saccadic reaction time. J Neurosci 31 (12), 4698–708.

71. Gaillard, C. et al. (2020) Prefrontal attentional saccades explore space rhythmically. Nat Commun 11 (1), 925.

72. Neupane, S. et al. (2017) Coherent alpha oscillations link current and future receptive fields during saccades. Proc Natl Acad Sci U S A 114 (29), E5979–E5985.

73. Zanos, T.P. et al. (2016) Mechanisms of Saccadic Suppression in Primate Cortical Area V4. J Neurosci 36 (35), 9227–39.

74. Payne, B.R. et al. (2019) Event-related brain potentials reveal how multiple aspects of semantic processing unfold across parafoveal and foveal vision during sentence reading. Psychophysiology 56 (10), e13432.

75. Stites, M.C. et al. (2017) Getting ahead of yourself: Parafoveal word expectancy modulates the N400 during sentence reading. Cogn Affect Behav Neurosci 17 (3), 475–490.

76. Cree, G.S. et al. (1999) An attractor model of lexical conceptual processing: Simulating semantic priming. Cognitive Science 23 (3), 371–414.

77. McLeod, P. et al. (2000) Attractor dynamics in word recognition: converging evidence from errors by normal subjects, dyslexic patients and a connectionist model. Cognition 74 (1), 91–113.

78. Devereux, B.J. et al. (2018) Integrated deep visual and semantic attractor neural networks predict fMRI pattern-information along the ventral object processing pathway. Sci Rep 8 (1), 10636.

79. Daelli, V. and Treves, A. (2010) Neural attractor dynamics in object recognition. Exp Brain Res 203 (2), 241–8.

80. Brunel, N. and Lavigne, F. (2009) Semantic priming in a cortical network model. J Cogn Neurosci 21 (12), 2300–19.

81. van Es, M.W.J. et al. (2020) Phasic modulation of visual representations during sustained attention. Eur J Neurosci.

82. Bahramisharif, A. et al. (2013) Propagating neocortical gamma bursts are coordinated by traveling alpha waves. J Neurosci 33 (48), 18849–54.

83. Wang, L. et al. (2018) Specific lexico-semantic predictions are associated with unique spatial and temporal patterns of neural activity. Elife 7.

84. Wang, L. et al. (2020) Neural Evidence for the Prediction of Animacy Features during Language Comprehension: Evidence from MEG and EEG Representational Similarity Analysis. J Neurosci 40 (16), 3278–3291.

85. Kuperberg, G.R. and Jaeger, T.F. (2016) What do we mean by prediction in language comprehension? Lang Cogn Neurosci 31 (1), 32–59.

